# Calcium-sensitive subthreshold oscillations and electrical coupling in principal cells of mouse dorsal cochlear nucleus

**DOI:** 10.1101/2023.07.27.550887

**Authors:** H Hong, LA Moore, PF Apostolides, LO Trussell

## Abstract

In higher sensory brain regions, slow oscillations (0.5-5 Hz) associated with quiet wakefulness and attention modulate multisensory integration, predictive coding, and perception. Although often assumed to originate via thalamocortical mechanisms, the extent to which sub-cortical sensory pathways are independently capable of slow oscillatory activity is unclear. We find that in the first station for auditory processing, the cochlear nucleus, fusiform cells from juvenile mice (of either sex) generate robust 1-2 Hz oscillations in membrane potential and exhibit electrical resonance. Such oscillations were absent prior to the onset of hearing, intrinsically generated by hyperpolarization-activated cyclic nucleotide-gated (HCN) and persistent Na^+^ conductances (NaP) interacting with passive membrane properties, and reflected the intrinsic resonance properties of fusiform cells. Cx36-containing gap junctions facilitated oscillation strength and promoted pairwise synchrony of oscillations between neighboring neurons. The strength of oscillations were strikingly sensitive to external Ca^2+^, disappearing at concentrations > 1.7 mM, due in part to the shunting effect of small-conductance calcium-activated potassium (SK) channels. This effect explains their apparent absence in previous *in vitro* studies of cochlear nucleus which routinely employed high-Ca^2+^ extracellular solution. In contrast, oscillations were amplified in reduced Ca^2+^ solutions, due to relief of suppression by Ca^2+^ of Na^+^ channel gating. Our results thus reveal mechanisms for synchronous oscillatory activity in auditory brainstem, suggesting that slow oscillations, and by extension their perceptual effects, may originate at the earliest stages of sensory processing.

**SIGNIFICANCE STATEMENT (120 words max.):** Many studies show that electrical activity in higher brain regions is regulated by brain oscillations. Here we show that such oscillatory activity can arise even in the first levels of auditory processing in the cochlear nucleus of the brainstem (fusiform cells), and is generated not by neural networks but by the biophysical properties of individual neurons. Oscillations are highly sensitive to external Ca^2+^ due to interplay of multiple ionic conductances. Gap junctions between cells allows for amplification and synchrony of such activity. Oscillations are absent in pre-hearing neurons, suggesting that sound activity might be important for their emergence. We propose that such early-level oscillations may serve to enhance signaling associated with particular environmental stimuli.

## INTRODUCTION

Oscillatory activity in the brain occurs at the single neuron and population level, typified by a rhythmic modulation of intracellular membrane potentials and extracellular local field potentials, respectively (Lagier et al., 2004; Okun et al., 2010; Hong et al., 2022). Distinct oscillation frequency bands seem to correlate with distinct brain states and arousal levels (Buzsáki, 2006), such that specific oscillation bands may promote coherent population activity, enable neurons to respond optimally to inputs of a certain frequency, and facilitate synaptic plasticity (Huerta and Lisman, 1995; Hutcheon and Yarom, 2000; Buzsáki and Draguhn, 2004; Lee et al., 2018).

In sensory cortices, slow oscillations in the 0.5-5 Hz band are particularly associated with deep sleep, quiet wakefulness (Crochet and Petersen, 2006; Poulet and Petersen, 2008) and correlate with stimulus familiarity (Kissinger et al., 2018) as well as perceptual decisions (Einstein et al., 2017). In the auditory system, these slow oscillations are proposed to modulate cellular and circuit excitability (Lakatos et al., 2005) and amplify sensory inputs (Lakatos et al., 2007). Indeed, recent studies in mice and humans suggest that the relative phase of slow oscillations modulates the neural discrimination and perceptual detection of sound features (Henry et al., 2016; Guo et al., 2017), with slow oscillations predictively entraining to subthreshold rhythmic sounds to enable faster perception (Lawrance et al., 2014; ten Oever et al., 2014, 2017). Altogether, these previous studies suggest that slow oscillations dynamically influence the excitability of higher-order sensory circuits, such that understanding the origin and mechanistic bases of slow oscillations could identify fundamental determinants of sensory perception.

Although not fully understood mechanistically, slow oscillations are generally assumed to originate via intracortical or thalamo-cortical interactions (Sanchez-Vives & McCormick, 2000; Beltramo et al., 2013; Neske, 2016). By contrast, whether slow oscillations can be independently generated by sensory circuits below the thalamus has received little attention. We recently showed that in auditory efferent neurons of the brainstem, intrinsic ion channel properties are sufficient to drive membrane oscillations that lead to infra-slow (∼0.1 Hz) bursting activity of the neuron (Hong et al., 2022). Here we show a different pattern of slow oscillatory activity (1-2 Hz) in the auditory brainstem that occurs in the first station of auditory afferent pathway. Using patch-clamp electrophysiology in acute brain slices of mouse auditory brainstem, we found that the principal output neurons (fusiform cells) of the dorsal cochlear nucleus (DCN) display large, spontaneous, and highly regular 1-2 Hz membrane potential oscillations. Such oscillations are absent before hearing onset. Pharmacology revealed that these oscillations are generated by the interplay of two ionic conductances, the hyperpolarization-activated cyclic nucleotide-gated channels (HCN) and tetrodotoxin-sensitive Na^+^ channels. In exploring why such oscillations have not been previously reported in brain slice studies, we found that they were acutely sensitive to external Ca^2+^ concentration over the range used in standard *in vitro* studies, and disappeared in concentrations greater than 1.7 mM. Finally, due to the electrical coupling between fusiform cells, subthreshold oscillations were strengthened and synchronized between nearby neurons, and this coupling and synchrony appeared to be confined to fusiform cells within single tonotopic domains. These results have significant implications for our understanding of the initiation and propagation of oscillatory activity throughout the central nervous system.

## MATERIALS AND METHODS

### Animals

All animal procedures were approved by Oregon Health & Science University’s Institutional Animal Care and Use Committee. Mice of postnatal (P) days 16-25 were used from WT *C57BL/6*, heterozygous GlyT2-EGFP (*FVB.Cg-Tg(Slc6a5-EGFP)13Uze/UzeBsiRbrc*, RRID:IMSR_RBRC04708; Zeilhofer et al. 2005), Thy1-YFP (*B6;CBA-Tg(Thy1-YFP)GJrs/GfngJ*, RRID:IMSR_JAX:014130; Feng et al. 2000), Vglut2-IRES-Cre (*Slc17a6tm2(cre)Lowl/J*, RRID: IMSR_JAX:016963; Vong et al., 2011), Ai9 tdTomato reporter (*B6.Cg-Gt(ROSA)26Sortm9(CAG-tdTomato)Hze/J*, RRID:IMSR_JAX:007909; Madisen et al., 2010), and connexin36 KO (Hormuzdi et al., 2001) of both sexes. To study the development of oscillations, P8-9 WT mice were used. Fusiform cells are easily identifiable in slice preparations by their morphology and input resistance, but the GlyT2-EGFP and Thy1-YFP genetic lines were used to more efficiently visualize deeper cells and distinguish them from other cell types, particularly in older mice. Specifically, the Thy1-YFP line labeled fusiform cells, while the GlyT2-EGFP line allowed us to avoid the extremely numerous glycinergic interneurons of DCN surrounding the fusiform cells. In some experiments, offspring of Vglut2-IRES-Cre and Ai9 tdTomato crosses were used. No differences were observed in the properties of fusiform cells among these different mouse lines.

### Brain-slice preparation

Mice were decapitated under deep isoflurane anesthesia. Coronal slices (300 µm for P16-25 mice, 210 µm for P8-9 mice) were cut on a vibratome (Leica VT1200S and Campden Instruments 7000smz-2). Slices containing DCN were cut at a 30° ventral-dorsal angle (to preserve electrical coupling of fusiform cells) in warm (∼35°C) artificial cerebrospinal fluid (ACSF). We were unsuccessful in finding electrically coupled fusiform cell pairs when slices were cut in the sagittal plane (n=11 dual recordings; data not shown). ACSF contained (in mM) 130 NaCl, 2.1 KCl, 1.2 KH_2_PO_4_, 1.7 CaCl_2_, 1 MgSO_4_, 20 NaHCO_3_, 3 Na-HEPES, 10-12 glucose, 2 Na-pyruvate, and 10 µM MK-801 and was bubbled with 5% CO_2_/95% O_2_ (300-310 mOsm). Slices recovered for 30 min at 34°C and were then kept at room temperature until use.

### Electrophysiology

Slices containing DCN were held in a chamber perfused with ACSF at a rate of 2-3 ml/min and heated to 30-33°C with an in-line heater. Neurons were visualized with Dodt contrast optics using 10X and 40X objectives on an upright microscope (Zeiss Axioskop2). A Multiclamp 700B amplifier was used to obtain whole-cell patch-clamp recordings from fusiform cells. Data were filtered at 10 kHz, digitized at 20-50 kHz with a Digidata 1322A, or at 100 kHz with a Digidata 1440A, and acquired with pClamp 10.4 software (Molecular Devices, RRID:SCR_011323). Except for the data in Figure 1D, oscillations were recorded in the presence of synaptic blockers to reduce signal variance and optimize cross-correlation analysis, including 5-10 µM NBQX, 10 µM MK-801, 0.5-1 µM strychnine, and 5-10 µM SR-95531.

**Figure 1.**
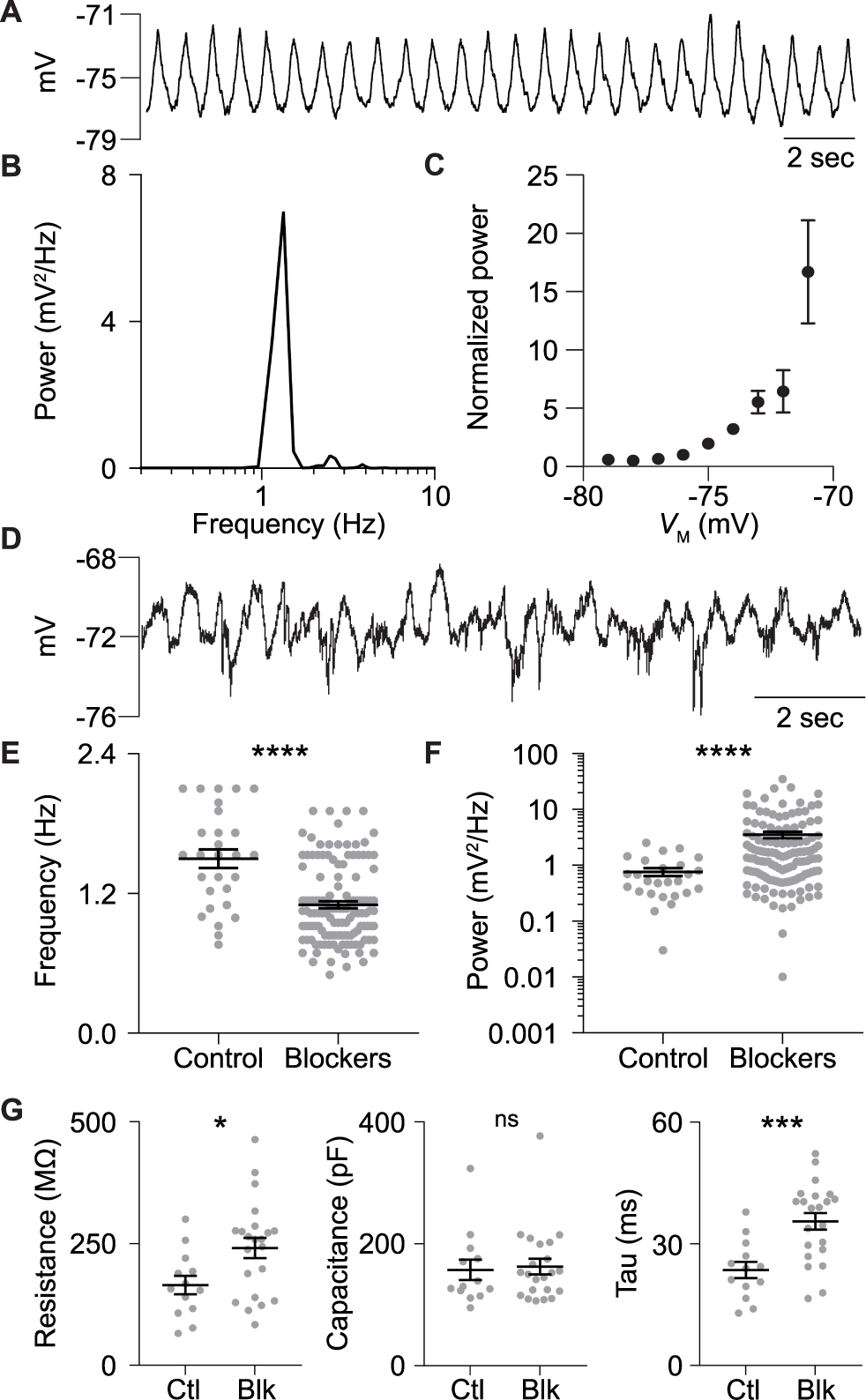
Fusiform cells display 1-2 Hz spontaneous oscillations of membrane potential. **A**, Example raw voltage trace from an oscillating fusiform cell in the presence of blockers of fast synaptic transmission (10 µM NBQX, 10 µM MK-801, 0.5 µM strychnine and 10 µM SR-95531). **B**, Power spectrum for example cell shown in **A**, with a peak between 1 and 2 Hz. **C**, Voltage dependence of subthreshold oscillations (n=10 cells, power is normalized to the power measured at -76 mV) in synaptic blockers. **D**, Example raw voltage trace showing oscillations in the absence of synaptic blockers. **E-F**, Population data showing oscillation frequency and power in control and with synaptic blockers. **G**, Population data showing the input resistance, membrane capacitance and time constant (tau) in control and with synaptic blockers. *p<0.05, ***p<0.001, ****p<0.0001. Error bar, SEM.

For experiments in which the bath Ca^2+^ concentration was varied, the total concentration of divalent ions was held constant at 2.7 mM by addition of Mg^2+^ to the ACSF. ACSF of each Ca^2+^ concentration contained (in mM) 130 NaCl, 3 Na-HEPES, 20 NaHCO_3_, ∼7 glucose, 0.05 NiCl_2_, 0.05 CdCl_2_, 0.4 ascorbate, and 2 Na-pyruvate. ACSF of varying Ca^2+^concentrations were mixed from stock solutions containing either 2.6 mM Mg^2+^ or 2.6 mM Ca^2+^. The Mg^2+^ stock solution contained, along with the above, 1 MgSO_4_, 1.6 MgCl_2_, and 3.3 KCl, and the Ca^2+^ stock solution contained 2.6 CaCl_2_, 1 K_2_SO_4_, and 1.3 KCl. KCl replaced KH_2_PO_4_ in order to prevent precipitation of Ca^2+^ (Kim and Trussell, 2007). HCN conductance in varying concentrations of Ca^2+^ was measured in a total divalent concentration of 2.8 mM in order to include 0.2 mM BaCl_2_.

The majority of experiments were performed with a K-gluconate internal pipette solution containing, in mM: 113 K-gluconate, 4.5 MgCl_2_, 0.1 EGTA, 9 HEPES, 14 tris-phosphocreatine, 4 Na_2_-ATP, 0.3 tris-GTP, with pH adjusted to 7.25 with KOH and osmolality adjusted to 290 mOsm with sucrose. In some experiments, 4.5 mM MgCl_2_ were replaced by 2.75 mM MgCl_2_ and 1.75 mM MgSO_4_. Membrane potentials measured using the K-gluconate pipette fill were adjusted for a junction potential of -13 mV. Input resistance was measured by injecting a -10-pA current into the fusiform cell in current-clamp mode. A Cs-TEA internal was used to measure persistent Na^+^ (NaP) conductance. This internal solution contained (in mM) 115 TEA-Cl, 4.5 MgCl_2_, 3.5 CsCl, 10 EGTA, 10 HEPES, 4 Na_2_-ATP, 0.5 tris-GTP, 5 CsOH to adjust pH to 7.25, and 15 sucrose to adjust osmolality to 290 mOsm. Results from experiments using this solution were corrected for a junction potential of -3 mV. Recording electrodes of 4-6 MΩ were pulled from borosilicate glass (WPI 1B150F-4) with a vertical puller (Narishige P-10). For voltage-clamp recordings, series resistance (<20 MΩ) was compensated by 60-80% correction and 90% prediction with the Multiclamp 700B (bandwidth 3 kHz). Data were only included for final analysis if the series resistance changed by <20% over the course of the experiment. Reagents were purchased from Sigma-Aldrich and Alomone Labs.

### Frequency-domain analysis

The power spectrum for spontaneous oscillations (binned at 0.1-0.2 Hz) was generated in Clampfit analysis software (version 10.9, Molecular Devices) from 20-30 s segments of representative traces using a Hamming window 50% overlap of spectral segments. The resonant frequency of neurons was determined by injecting a 5 pA swept sine wave (ZAP, or impedance (*Z*) amplitude profile) current that varied in frequency from 0 to 10 Hz over 20 seconds (Puil et al., 1986). Neurons were injected with negative bias current just sufficient to prevent spiking during the ZAP current injection and spontaneous oscillation frequency and power were measured at this same holding potential. ZAP current was run for 20-50 trials for individual neurons and the voltage responses were averaged. Based on code from Bakken et al., (2021), impedance (*Z*) was calculated from the ratio of the fast Fourier transforms of measured voltage response (*V*) and the injected current (*I*):

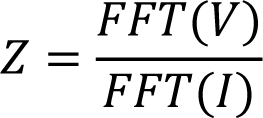

For a resonant neuron, the resulting impedance curve displays a peak in impedance at the resonant frequency. A smoothing protocol was applied to reduce flutuations in the impedance curve (Fig. 4B), and measurements of resonance peak made from the smoothed trace. The height of this peak is quantified by the Q value, which here is the peak impedance value divided by the impedance value measured at 0.4 Hz.

### Conductance measurements

To measure NaP activation, a ramp was delivered to the neuron from -83 to -53 mV over the course of 5 seconds (Leao et al., 2012). 1-10 nM TTX was included in the bath to help suppress escaping spikes and therefore enable current measurements at more depolarized potentials. Cd^2+^/Ni^2+^ (50 µM each) were included in the bath to block calcium channels. The first ∼2 seconds of the response was dominated by a linear outward current reflecting the passive membrane response to the voltage ramp. We fit this with a line and subtracted the linear values from the current response to perform leak subtraction. In a subset of experiments, a ramp from -83 to -3 mV over the course of 6 seconds was applied to the neuron. After recording the current response to this ramp, TTX (1 µM) was applied in the bath and the ramp was repeated. The current traces after TTX were subtracted from those before TTX, in order to isolate TTX-sensitive NaP current. These experiments were conducted with either 1.7 or 2.5 mM Ca^2+^. The results from the two voltage ramp protocols were consistent (see *RESULTS*). To analyze the voltage dependence of NaP activation, conductance (*G*) was calculated according to the following equation, where *I* is current measured at a given voltage (*V*), and *Vrev* is the reversal potential of Na^+^ (calculated to be +77.4 mV):

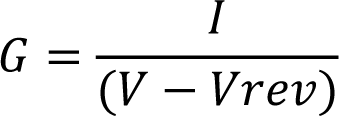

HCN activation in different external Ca^2+^ concentrations was measured similar to Tang & Trussell (2015). Cells were held at -60 mV and 5-second steps from -120 to -60 mV were applied in 5-mV increments. After each 5-second step, the cell was brought back to -60 mV, resulting in a tail current from which HCN activation could be approximated. The bath solution included Cd^2+^/Ni^2+^ (50 µM each) to block Ca^2+^ currents, and TTX (0.5 µM) to block Na^+^ current. Ba^2+^ (0.2 mM) and 4-aminopyridine (4-AP; 1 mM) were also included to reduce contamination in the tail current from leak conductance and low-voltage activated K^+^ channels respectively (Johnston et al., 2010; Leao et al., 2012). Both drugs slightly block HCN channels at these concentrations (Ludwig et al., 1998).

The resulting NaP and HCN activation curves were fit with a Boltzmann equation:

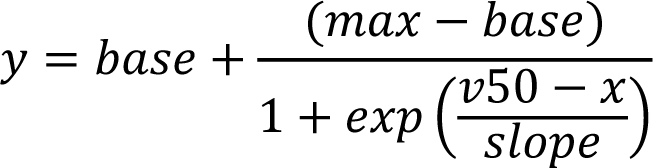

The *base* value was set to 0. The other resulting values, *max* (maximum conductance), *v50* (half-activation voltage of the curve), and *slope* (a value that describes the steepness of the sigmoid) were compared for each cell between different external Ca^2+^ concentrations. The population conductance curves for NaP and HCN were generated by normalizing raw conductance values for each cell to the *max* value given by the Boltzmann fit. These average values were then fit with their own Boltzmann equation.

### Paired recordings and cross-correlation analysis

The coupling coefficient for electrically coupled fusiform cells was determined by holding both cells in current clamp and alternating injection of a -1 nA square pulse into each cell. Coupling coefficient was calculated by dividing the amplitude of the voltage deflection in the post-junctional cell by the amplitude of the voltage deflection in the pre-junctional cell. Pairs were considered to be coupled if the coupling coefficient was greater than 0.005 (Yaeger and Trussell, 2016).

For cross-correlation analysis for cell pairs, 1-2 min segments of subthreshold traces with a steady baseline were selected. Traces were resampled from 50 to 200 µs inter-sample-interval, bandpass filtered at 0.2-10 Hz, and baseline-subtracted. Cross-correlograms were generated in Igor and normalized by dividing cross-correlation coefficient values (xcorr) by the standard deviation (sd) of each trace multiplied together:

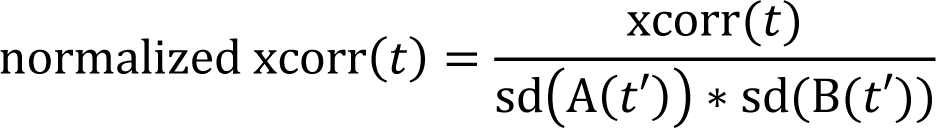

where xcorr(t) is the raw cross-correlation coefficient calculated for a given lag time t, and A(t’) and B(t’) are the voltage traces with t’ representing the time variable for the traces. In this way, if the analysis were an auto-correlation, variance would be the normalization factor and the value at time zero would be 1 (Beierlein et al., 2000).

For spike cross-correlation, spikes were detected using threshold search in Clampfit, generating a list of spike times for each cell. Spike times were binned in 1- and 10-ms segments and cross-correlated in Igor. Resulting cross-correlation coefficients (xcorr) were then adjusted and normalized according to the following equation:

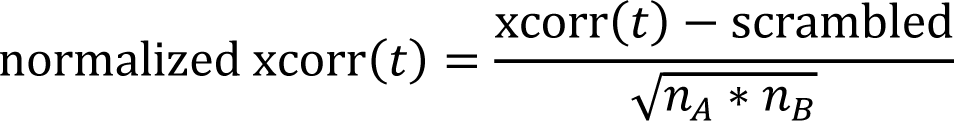

where n_A_ and n_B_ are spike counts from traces A and B. The *scrambled* term is the average cross-correlation coefficient value from cross correlating spikes from trace A with scrambled spike times from trace B, generated using the *random.shuffle* function in Python. This is used in place of the theoretically calculated term used by others (Eggermont, 1992; Stefanescu and Shore, 2015). In the text, both oscillation and spike time cross-correlation coefficient values are referred to as *xcorr* for simplicity. Peak *xcorr* values reported are measured at zero lag instead of the real peak.

### Experimental design and statistical analysis

Electrophysiological traces were analyzed with pClamp 10.4 software or custom-written procedures in IGOR Pro 6.3 or 8.04 (RRID:SCR_000325) and Python 2.6 (RRID:SCR_008394). For spike shape analysis in Extended Data Table 2-1, spike threshold was defined as voltage at the time point when spike dV/dt reached 5% of its peak. GraphPad Prism 7 and 9 (RRID:SCR_002798) was used for statistical analysis and to make figures. Data are represented in the text and figures as mean ± SEM. Parametric analysis was used only after confirming assumptions of equal variances (F test for two-group comparison and Bartlett’s test for three-group comparison) and normality (D’Agostino-Pearson omnibus normality test). The equivalent non-parametric tests were used when data were not normal; the name of the test used is included with all statistics reported in the *RESULTS* section. One-way ANOVA with repeated measures (RM) was used to compare statistics for single cells across more than two conditions, followed by Tukey’s *post hoc* comparisons (p values associated with post hoc tests are only reported when the main effect is significant). The Geisser and Greenhouse correction was applied to RM one-way ANOVAs to correct for possible violations of the assumption of sphericity (of note, this frequently changes associated degrees of freedom values to non-integers). Assumptions of sphericity and effective matching were also confirmed for one-way ANOVAs.

## RESULTS

### Fusiform cell membrane potential oscillates at rest

In acute DCN brain slices, approximately 80% of patched fusiform cells displayed 1-2 Hz subthreshold oscillations around the resting membrane potential in the absence of ongoing spike activity (Fig. 1A). Power spectrum analysis of the oscillations revealed a clear peak in this frequency range, averaging 1.10 ± 0.03 Hz over the population (n=128 cells; Fig. 1B,E). Oscillation power was strongly voltage-dependent, indicating the involvement of voltage-sensitive channels (Fig. 1C; n=10 cells). Oscillations were typically recorded in the presence of blockers of fast synaptic transmission (see *MATERIALS AND METHODS*), but were also clearly present in the absence of synaptic blockers (Fig. 1D). Synaptic blockers slightly reduced the oscillation frequency (Fig. 1E; control: 1.50 ± 0.08 Hz, n=28 cells; synaptic blockers: 1.10 ± 0.03 Hz, n=128 cells; U=845, p<0.0001, Mann-Whitney U) but increased the oscillation power (Fig. 1F; control: 0.8 ± 0.1 mV^2^/Hz, n=26 cells; synaptic blockers: 3.5 ± 0.5 mV^2^/Hz, n=128 cells; U=862, p<0.0001, Mann-Whitney U). The increase in power with blockers was paralleled by an increase in input resistance (Fig. 1G, left; control: 164.8 ± 18.8 MΩ, n=13 cells; synaptic blockers: 240.9 ± 20.8 MΩ, n=22 cells; t(33)=2.478, p=0.019, two-tailed t-test), reflecting the loss of the shunting effect of spontaneous synaptic activity. Accordingly, membrane time constant (tau) increased upon block of synaptic activity (Fig. 1G, right; control: 23.56 ± 2.02 ms; synaptic blockers: 35.54 ± 2.06 ms; t(33)=3.860, p=0.0005, two-tailed t-test), while membrane capacitance did not change (Fig. 1G, middle; U=135, p=0.8, Mann-Whitney U).

### Oscillations are absent in fusiform cells from pre-hearing mice

We next asked whether the presence of oscillations reflected an electrically immature state of fusiform cells by recording from younger neurons, specifially choosing ages (P8-9) before hearing onset. None of the fusiform cells generated spontaneous action potentials (APs) at these immature ages. Unlike older neurons, the majority of P8-9 fusiform cells (12 out of 13) did not exhibit oscillations at rest (Fig. 2A). The power from auto-correlation analysis was below 0.1 mV^2^/Hz for all the pre-hearing neurons, significantly smaller than that of P16-25 fusiform cells (Fig. 2B; P8-9: 0.030 ± 0.007 mV^2^/Hz; n=13 cells, U=11, p<0.0001, Mann-Whitney U). Surprisingly, such profound changes in oscillation did not correlate with developmental changes in passive membrane properties, as assessed with small hyperpolarizing steps (Fig. 2C; input resistance: P8-9: 269.7 ± 17.87 MΩ, P16-21: 240.9 ± 20.75 MΩ, U=126, p=0.38; capacitance: P8-9: 178.0 ± 12.75 pF, P16-21: 162.5 ± 12.84 pF, U=111, p=0.17; tau: P8-9: 46.44 ± 2.82 ms, P16-21: 35.54 ± 2.06 ms, U=76.5, p=0.01, Mann-Whitney U). Besides oscillations, active properties also changed with age. In 11 out 14 fusiform cells at P8-9, a delay to onset of spiking was apparent, an effect often attributed to activation of A-type K^+^ current (Fig. 2D, arrowhead, Kanold and Manis, 1999). This delay was not observed at older ages (Fig. 2E). Fusiform cells from these two age groups also showed significant differences in rheobase, AP threshold and AP width (Extended Data, Table 2-1), suggesting changes in Na^+^-channel expression. Taken together, the parallel maturation of AP properties and oscillation strength with age suggests that some developmentally regulated ion channels play a role in both.

**Figure 2.**
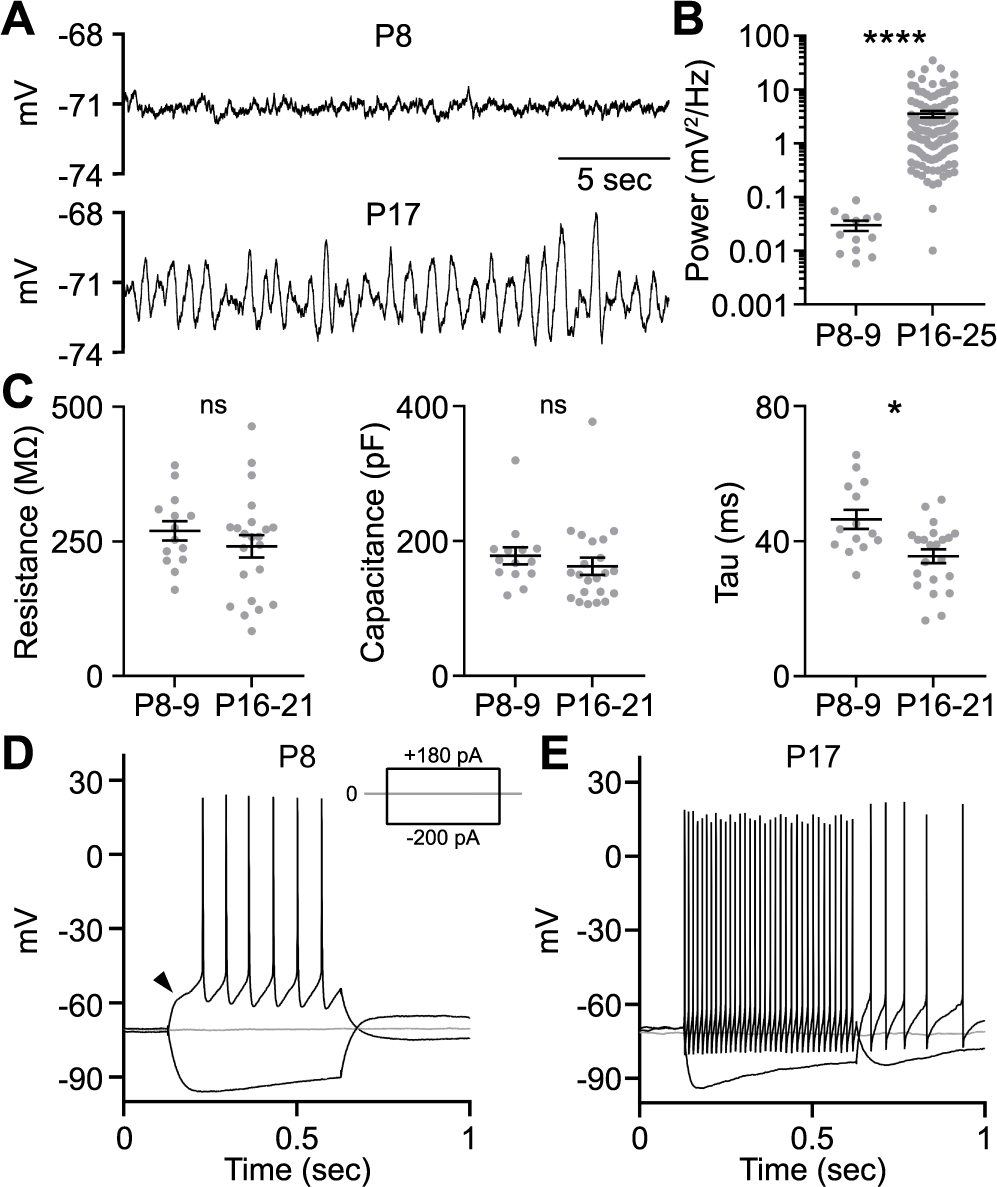
Oscillations are absent in P8-9 fusiform cells. **A**, Representative voltage traces from fusiform cells at P8 (before hearing onset) and P17 (after hearing onset). **B**, Population data showing the oscillation power in two age groups. Note that P16-25 data are the same as Figure 1F, right column. **C**, Population data showing input resistance, membrane capacitance and time constant (tau) in two age groups. **D-E**, Raw voltage traces from the two representative neurons shown in **A** in response to current injections at -200, 0 and +180 pA. Arrowhead points to the delay before the first action potential. *p<0.05, ****p<0.0001. Error bar, SEM. Recordings made in the presence of blockers of fast synaptic transmission (5-10 µM NBQX, 10 µM MK-801, 0.5-1 µM strychnine and 5-10 µM SR-95531).

### NaP and HCN conductances are required for oscillations

Previous studies show that subthreshold oscillations can be generated by Na^+^ channels (i.e., NaP current), HCN and/or T-type Ca^2+^ channels (Mccormick and Pape, 1990; Hughes et al., 2002; Fransén et al., 2004). We therefore explored whether these ion channels participate in driving the oscillations in fusiform cells. First, addition of 0.5 µM TTX markedly reduced oscillations, indicating a critical role for Na^+^ channels (Fig. 3A; control: 6.1 ± 4.3 mV^2^/Hz; TTX: 0.0052 ± 0.0009 mV^2^/Hz; wash: 4.6 ± 3.4 mV^2^/Hz; n=7 cells, W=11.14, p=0.0012, Friedman test). Overall, 88% (22 out of 25) of fusiform cells recorded in TTX showed no oscillations. Second, bath application of the HCN channel blocker Cs^+^ (1-2 mM) also attenuated oscillation power (Fig. 3B; control: 5.7 ± 1.8 mV^2^/Hz; Cs^+^: 0.2 ± 0.1 mV^2^/Hz; wash: 4.6 ± 1.4 mV^2^/Hz; n=10 cells, F(1.2, 10.8)=8.7, p=0.01, RM one-way ANOVA). In a separate set of experiments, ZD7288 (10 μM) was applied to confirm the role of HCN channels in oscillations. ZD7288 had matching effects to Cs^+^ (control: 8.0 ± 3.9 mV^2^/Hz; ZD: 0.4 ± 0.03 mV^2^/Hz; n=6 cells, W=-21, p=0.03, Friedman test).

**Figure 3.**
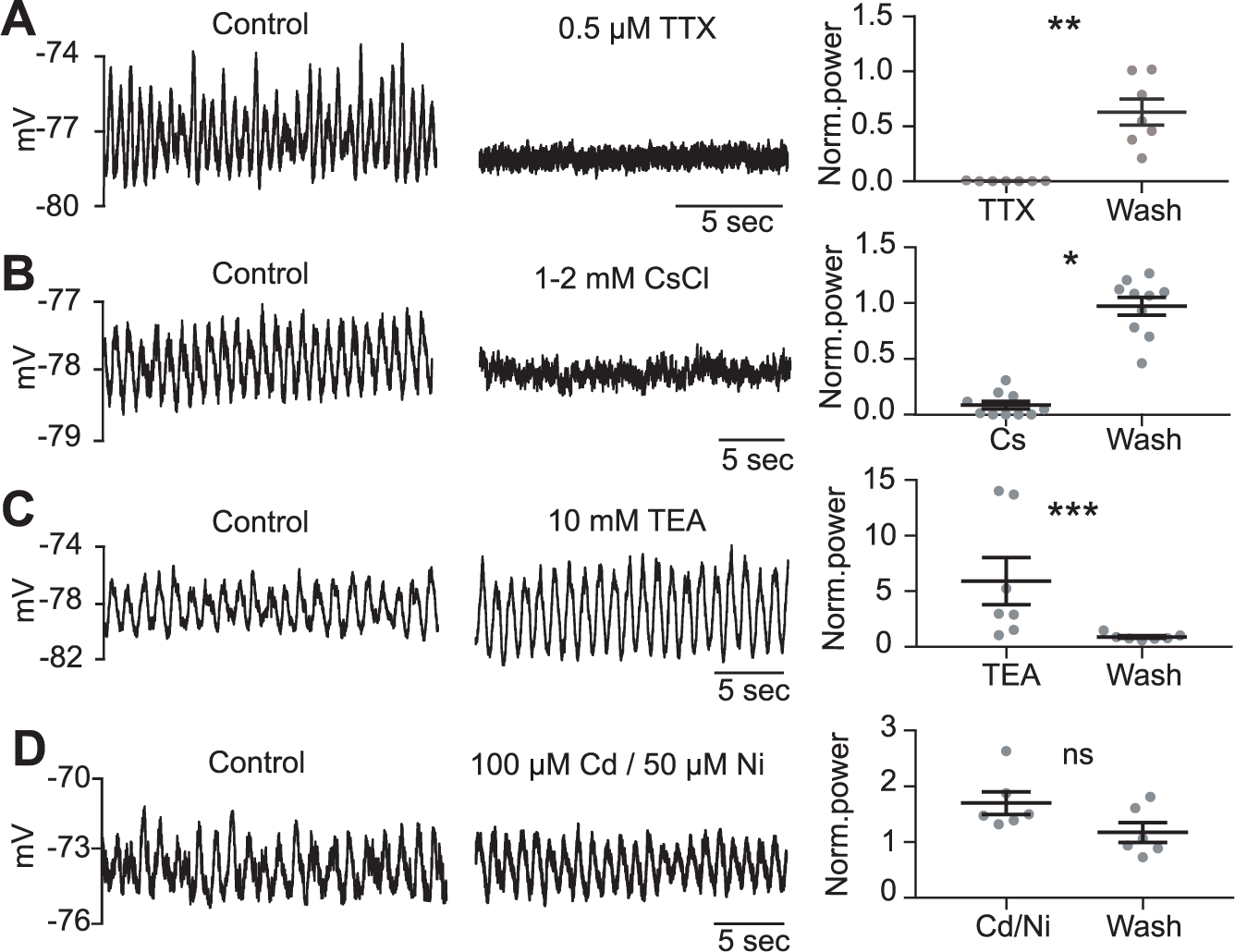
Subthreshold oscillations require persistent Na^+^ (NaP) and HCN conductances. Oscillations were blocked by bath application of 0.5 µM TTX (**A**; W=11.14, p=0.0012, Friedman test) and 1-2 mM CsCl (**B**; F(1.2, 10.8)=8.7, p=0.01, RM one-way ANOVA). Conversely, 10 mM TEA (**C**) increased oscillation power (F=12.29, p=0.0003, Friedman test). Broad-spectrum Ca^2+^ channel blockers (**D**) also appeared to increase oscillations, but the enhancement was not significant (F(1.6,7.9)=4.64, p=0.052, RM one-way ANOVA). Bias current was applied to correct for membrane potential changes with drug application before power measurements were made. Population graphs are normalized to control. *p<0.05, **p<0.01, ***p<0.001. Error bar, SEM. Recordings made in the presence of blockers of fast synaptic transmission (5-10 µM NBQX, 10 µM MK-801, 0.5-1 µM strychnine and 5-10 µM SR-95531).

We also tested for the involvement of K^+^ channels. The non-specific K^+^ channel blocker TEA (10 mM) significantly increased oscillation power from 2.5 ± 1.9 to 13.7 ± 9.8 mV^2^/Hz (Fig. 3C; wash: 2.0 ± 1.5 mV^2^/Hz; n=7 cells, F=12.29, p=0.0003, Friedman test). This observation suggests that K^+^ channels activating at subthreshold voltages exert a shunting effect on membrane oscillations, rather than driving the repolarization of the oscillatory cycle. Finally, we tested the role of T-type Ca^2+^ channels, which are known to drive oscillatory activity in thalamus (Steriade et al., 1993; Hughes et al., 2002). Surprisingly, we found that oscillations in fusiform cells persisted in the presence of Ca^2+^ channel blockers: addition of broad-spectrum Ca^2+^ channel blockers, 100 μM Cd^2+^ and 50 μM Ni^2+^, resulted in a slight but non-significant increase in oscillation power (Fig. 3D; control: 3.9 ± 1.5 mV^2^/Hz; Cd^2+^/Ni^2+^: 7.2 ± 2.6 mV^2^/Hz; wash: 4.6 ± 1.6 mV^2^/Hz; n=6 cells, F(1.6,7.9)=4.64, p=0.052, RM one-way ANOVA). Ni^2+^ also blocks NMDA receptors and therefore rules out the involvement of this conductance as well (Mayer and Westbrook, 1985); the exposure of tissue to MK-801 during slice preparation further reduces the likelihood of NMDA receptor involvement. Nevertheless, the slight increase in oscillation power with Cd^2+^ and Ni^2+^ might be a result of indirect blockade of Ca^2+^-activated K^+^ channels, thus suggesting the possibility of a tonic activation of Ca^2+^-activated K^+^ channels at resting voltages. In a later section (Fig. 6), we demonstrate that this Ca^2+^-activated K^+^ channel is the small-conductance calcium-activated potassium (SK) channel. Altogether, these results support a model whereby oscillations are generated by the interplay between NaP and HCN conductances, whereas K^+^ channels negatively regulate the power of the oscillations.

### Fusiform cells show slow intrinsic resonance

Resonance refers to the ability of a neuron to amplify voltage responses to current stimuli over a narrow frequency range. To evaluate resonance, a frequency ramp (ZAP) is delivered to the neuron and the frequency-dependent resistance (impedance, *Z*) is measured across frequency (Hutcheon and Yarom, 2000); peak resonance corresponds to the peak in impedance. Previous studies show a close association between oscillations and resonance, as they may share underlying ion channel mechanisms (Hutcheon and Yarom, 2000). We explored resonance in fusiform cells by injecting a ZAP from 0-10 Hz (Fig. 4A). The raw impedance trace (Fig. 4B, grey trace) was smoothed (black trace) to identify the peak in the curve (dashed line, see *MATERIALS AND METHODS*). All the fusiform cells exhibited a resonant peak in response to the ZAP (n=19 cells; Fig. 4B). Notably, the frequency corresponding to the resonance peak was not significantly different from their oscillation frequency (Fig. 4C; resonance: 1.21 ± 0.09 Hz; oscillation: 1.23 ± 0.07 Hz; n=17 cells, t(16)=0.340, p=0.74, paired t-test). To characterize the magnitude of resonance, impedance at the resonance peak was normalized to impedance at 0.4 Hz (Fig. 4D). The peak of the resulting curve is defined as Q (Fig. 4E). A non-resonant neuron will have a Q value equal to 1. All fusiform cells measured had a Q value greater than 1, with an average value of 1.55 ± 0.06 (n=17 cells, U=0, p<0.0001, Mann-Whitney U).

**Figure 4.**
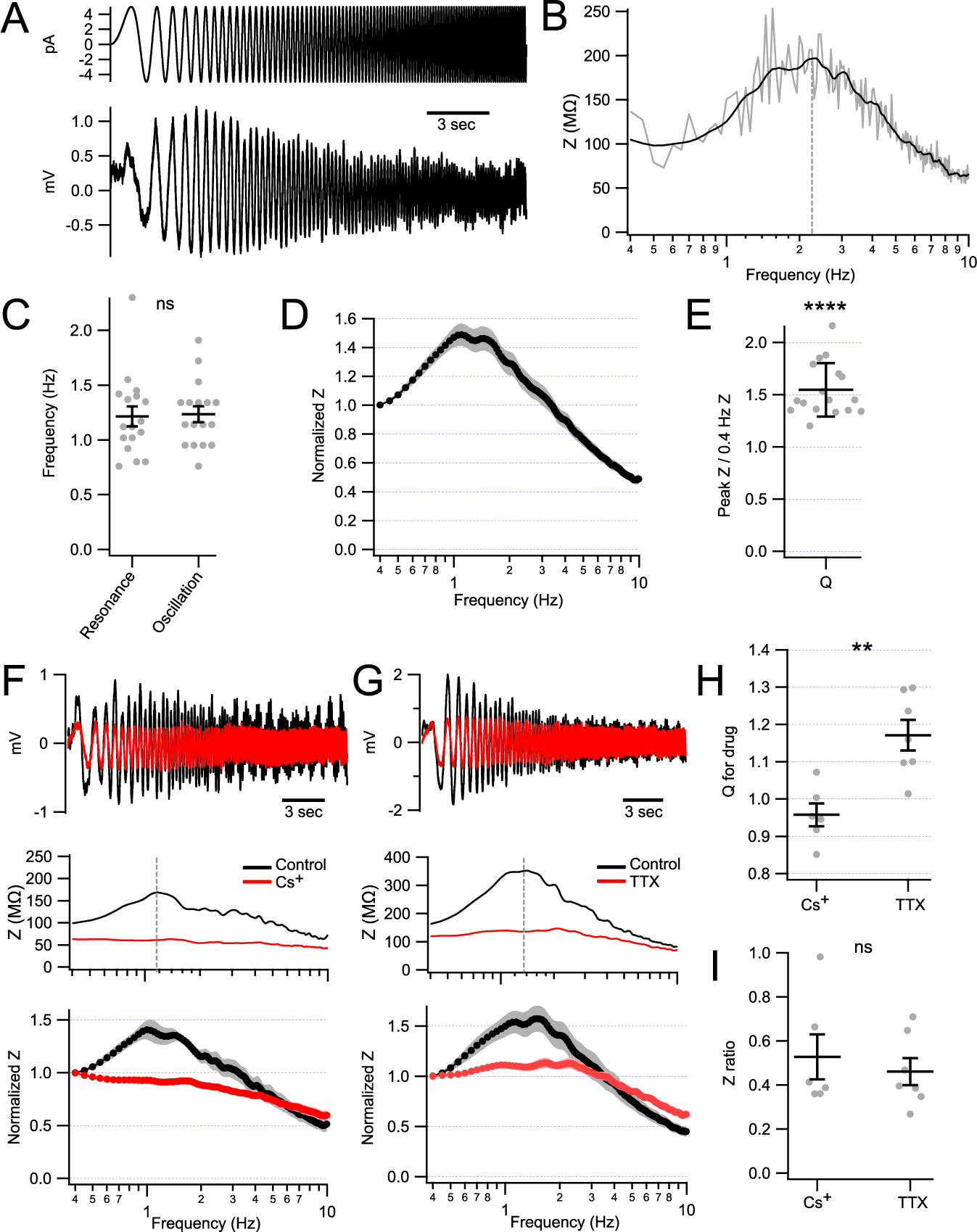
Close association between oscillation and membrane resonance. **A**, 5-pA ZAP current (upper panel) elicited resonant voltage response (averaged, lower panel) from a representative fusiform cell. **B**, Impedance (*Z*) from the cell shown in **A** plotted as a function of frequency. Grey and black traces represent impedance curve before and after smoothing, respectively. Vertical dashed line aligned at the peak of the curve. **C**, The resonance frequency of oscillating cells matched the frequency of their spontaneous oscillations (t(16)=0.340, p=0.739, paired t-test). **D,** Normalized impedance curve to *Z* at 0.4 Hz. The peak of this curve is defined as Q. **E,** Population data showing Q values of the resonance. ****p<0.0001 as compared to 1. **F-G,** Upper panel: representative voltage responses in control and after bath application of Cs^+^ (1 mM, **F**) or TTX (1 µM, **G**). Middle panel: raw impedance curve for the voltage response shown above. Lower panel: population data showing the normalized impedance curve. Vertical dashed line aligned at the peak of the curve. **H,** Population data showing Q for Cs^+^ or TTX. Q for drug was measured as the normalized impedance in the drug, measured at the frequency where Q occurred in control solutions. **p<0.01. **I,** Population data showing the *Z* ratio, calculated as remaining *Z* with Cs^+^ or TTX (measured at resonance frequency determined in control) divided by peak *Z* in control. Error bar, SEM. Recordings made in the presence of blockers of fast synaptic transmission (5-10 µM NBQX, 10 µM MK-801, 0.5-1 µM strychnine and 5-10 µM SR-95531).

As NaP and HCN conductances play an essential role in generating oscillations in fusiform cells, we next examined their roles in resonance. 1 mM Cs^+^ abolished resonance in fusiform cells (Fig. 4F), indicating a requirement for HCN channels in setting the resonance peak. The resulting impedance curve decreased as the frequency increased, a characteristic of a passive membrane (Fig. 4F, lower panel). For normalized impedance curves in the presence of Cs^+^ for each cell, we measured impedance value at the frequency corresponding to the resonance peak in control solutions. The average of this Q-for-drug value was 0.96 ± 0.03, not significantly different from 1 (Fig. 4H, left; n=6 cells, U=12, p=0.364, Mann-Whitney U). In addition, a “Z ratio” was calculated as the un-normalized impedance with Cs^+^ (again, measured at the resonance frequency determined in control) divided by the peak impedance in control, a metric that reflects both the changes in resonance and overall impedance induced by Cs^+^. On average, Z ratio was 0.53 ± 0.10 with Cs^+^ (Fig. 4I, left; n=6 cells).

TTX (0.5-1 µM) also attenuated the resonance of fusiform cells (Fig. 4G). Q in TTX was significantly smaller than control values (Fig. 4H, right; control: 1.70 ± 0.11; TTX: 1.17 ± 0.04; n=7 cells, t(6)=5.796, p=0.0012, paired t-test). Z ratio was 0.46 ± 0.06 in TTX (Fig. 4I, right). Different from what was observed with Cs^+^, however, neurons in TTX still maintained some residual resonance (Fig. 4G, lower panel), as the Q for TTX was significantly larger than 1 (U=0, p=0.0006, Mann-Whitney U), and larger than Q for Cs^+^ (Fig. 4H; t(11)=4.049, p=0.0019, two-tailed t-test). Taken together, HCN current is essential for resonance in fusiform cells, while NaP contributes to its amplification. These results show that oscillations are an outcome of the intrinsic resonance properties of fusiform cells generated by their complement of ion channels.

### External Ca**^2+^** concentration determines the strength of oscillations

The presence of oscillations in fusiform cells was surprising given that they had not been reported, despite decades of studies of these cells *in vitro* (Manis, 1990; Gardner et al., 2001; Fujino and Oertel, 2003; Manis et al., 2003; Tzounopoulos et al., 2004). However, most previous studies of fusiform cells used an external Ca^2+^ concentration of 2.4-2.5 mM (Gardner et al., 2001; Fujino and Oertel, 2003; Manis et al., 2003; Tzounopoulos et al., 2004). Our experiments thus far used 1.7 mM Ca^2+^, closer to the external Ca^2+^ concentration of CSF *in vivo* (Somjen 2004; Borst, 2010). Might differences in external Ca^2+^ explain why other studies did not report oscillations in fusiform cells? We tested this hypothesis by systematically determining the relation between oscillation power and Ca^2+^. Oscillation strength showed a remarkably steep dose-dependence on Ca^2+^ concentration, decreasing to undetectable levels above 2 mM Ca^2+^, with a half-maximal power at 1 mM (Fig. 5A-C, n=8 cells). Though 1.7 and 2.5 mM are on the tail-end of the dose-response curve, there is a >4 fold difference between power measurements in these two concentrations. Indeed, both raw and normalized power measurements show that power measured at 1.7 mM was significantly larger than power measured at 2.5 mM Ca^2+^ (U=-36, p=0.008, Mann-Whitney U, also see Fig. 6E). Figure 5C shows that oscillation power of a representative fusiform cell changed over time when external Ca^2+^ was switched between 1.7 and 0.5 mM. The interval between two consecutive data points is 30 seconds. The effect of Ca^2+^ was rapid: within three minutes of washing in 0.5 mM Ca^2+^, there was a pronounced increase in power, while the power quickly returned to its initial value when 1.7 mM Ca^2+^ was re-applied. Such increase in oscillation power is not due to altered input resistance, which was not significantly different between 1.7 and 0.5 mM Ca^2+^ (average(1.7 and 0.5 mM Ca^2+^)=114.3 ± 11.8, 115.4 ± 17.0 MΩ, n=5 cells, t(4)=0.082, p=0.939, paired t-test).

**Figure 5.**
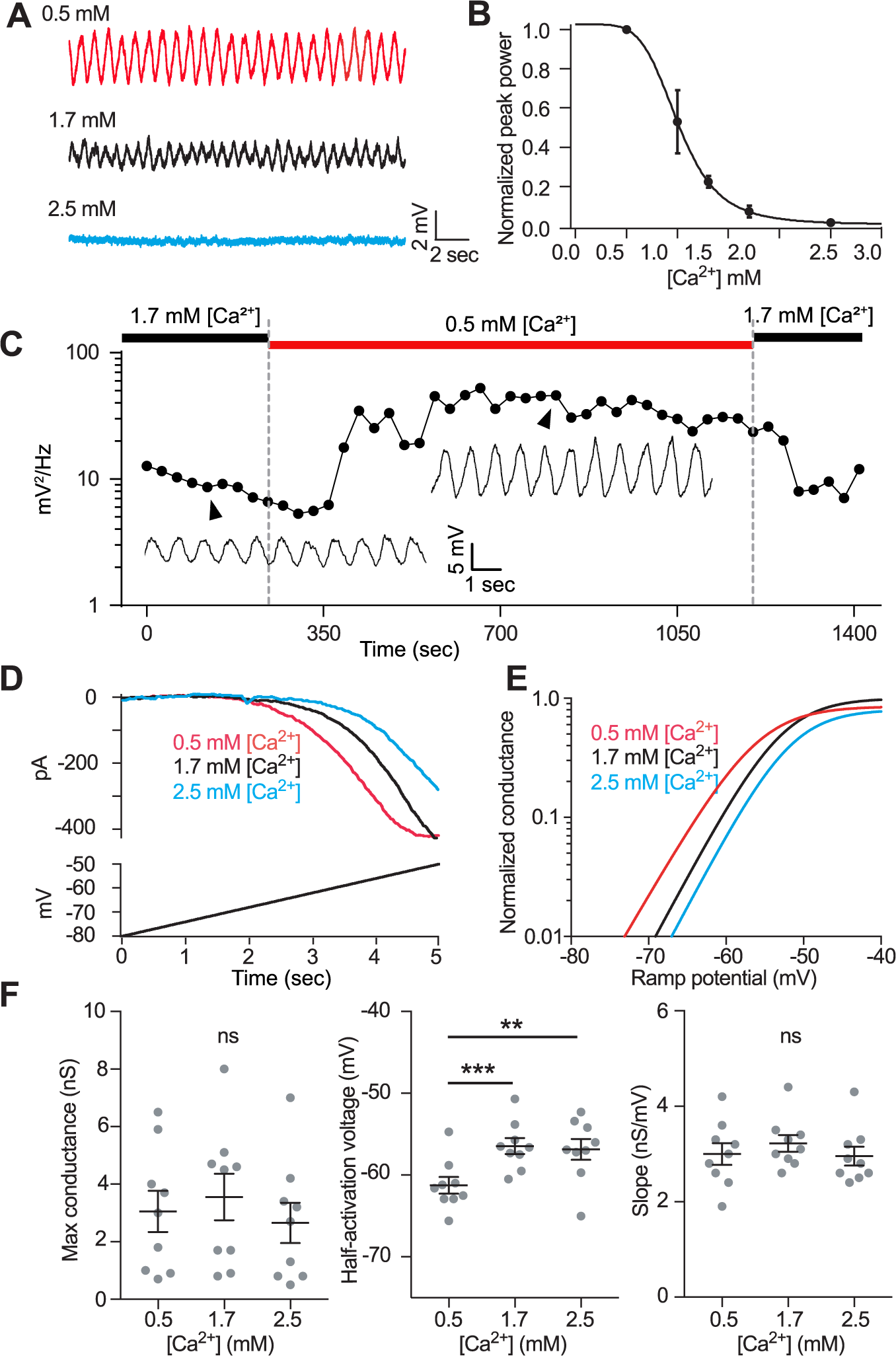
Lowering extracellular Ca^2+^ amplifies oscillations. **A**, Example raw voltage traces in different external [Ca^2+^]. **B**, Dose-response curve displaying oscillation power measured in fusiform cells across varying external [Ca^2+^] (n=8 cells). **C,** Power of oscillations from a representative fusiform cell plotted as a function of time. Arrowheads point to the time when the oscillations shown in the insets occurred. Note that insets are plotted in the same scale. **D**, Na^+^ conductance curves were measured in varying external [Ca^2+^] by applying a 5-second ramp holding from -83 to -53 mV. Raw current traces in response to the voltage ramp are displayed for one cell. Boltzmann curves from fits to the data in **D. F**, Population data comparing maximum conductance, half-activation voltage, and slope measured from Boltzmann fits across [Ca^2+^]. Maximum conductance was unchanged between condi..ons (F(1.2, 9.9)=1.6, p=0.239, RM one-way ANOVA). Half-activation voltage was positively shifted in [Ca^2+^] greater than 0.5 (F(1.7, 13.3)=31.07, p<0.0001, RM one-way ANOVA). There was no significant difference in slope between different external [Ca2^+^] (F(1.1, 9.0)=0.514, p=0.514, RM one-way ANOVA). *p<0.05, **p<0.01, ***p<0.001. Error bar, SEM. Recordings made in the presence of blockers of fast synaptic transmission (5-10 μM NBQX, 10 μM MK-801, 0.5-1 μM strychnine and 5-10 μM SR-95531).

**Figure 6.**
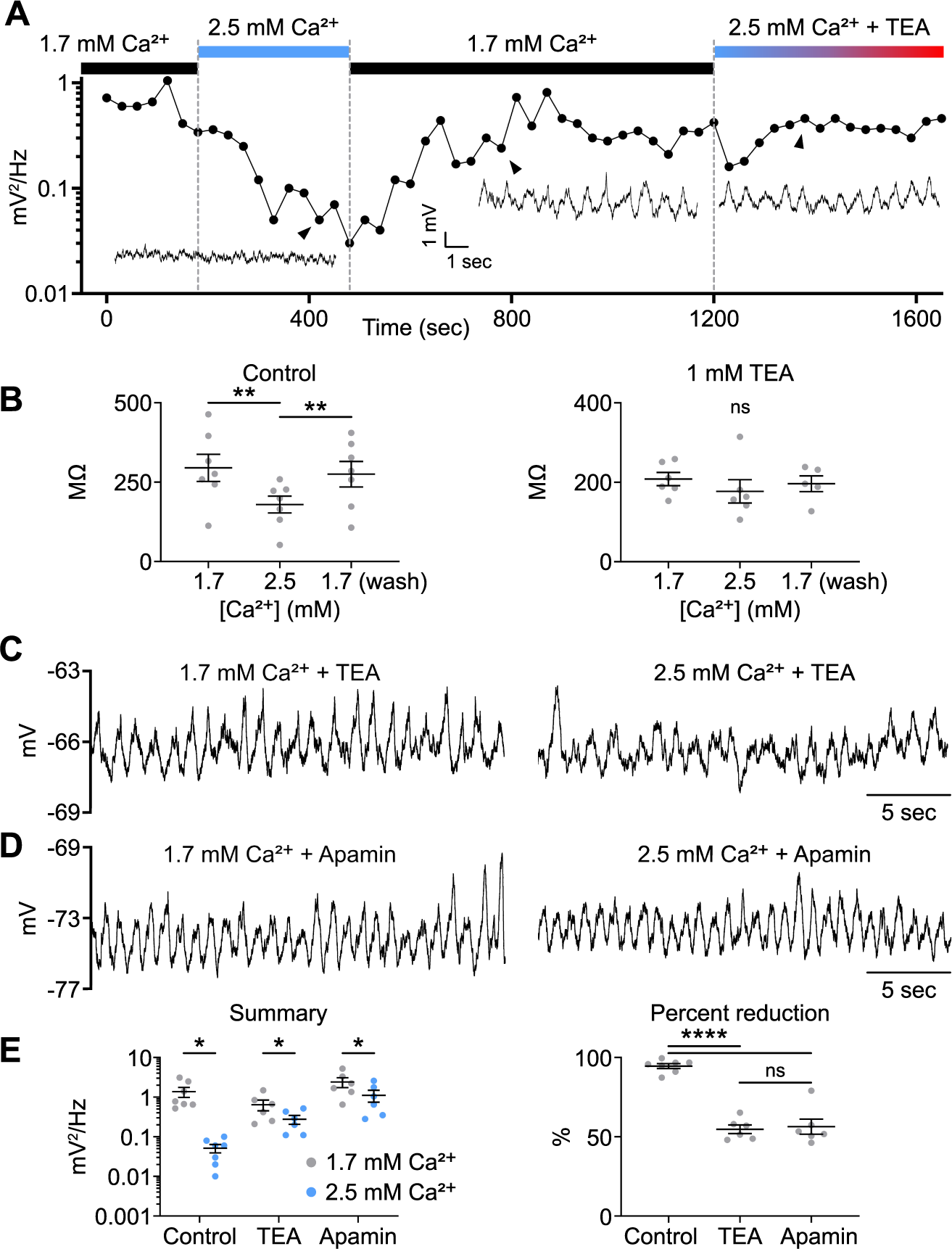
Increasing extracellular Ca^2+^ concentration activates SK channels that shunt oscillations. **A,** Power of oscillations from a representative fusiform cell plotted as a function of time. Arrowheads point to the time when the oscillations shown in the insets occurred. Note that insets are plotted in the same scale. **B,** Population data showing that the Ca^2+^ dependence of input resistance (left, q(6)=6.997, p=0.006, Tukey’s post hoc test) was abolished by 1 mM TEA (right, q(5)=2.121, p=0.367, Tukey’s post hoc test). **C-D,** Representative voltage traces recorded with 1 mM TEA (**C**) or 200 nM apamin (**D**) in the bath. **E,** Population data showing that the Ca^2+^ dependence of power was attenuated by application of TEA or apamin. Percent reduction of power: control *vs.* TEA, q(16)=12.80, p<0.0001; control *vs.* apamin, q(16)=12.28, p<0.0001, Tukey’s post hoc test. *p<0.05, **p<0.01, ****p<0.0001. Error bar, SEM. Recordings made in the presence of blockers of fast synaptic transmission (5-10 µM NBQX, 10 µM MK-801, 0.5-1 µM strychnine and 5-10 µM SR-95531).

The experiments described in Figure 3 highlight the critical role of Na^+^ and HCN channels in the genesis of the oscillations. Interestingly, previous studies showed that external Ca^2+^ concentration regulates the voltage sensitivity of Na^+^ channels, with lower concentrations causing a negative shift in the half-activation voltage (Frankenhaeuser and Hodgkin, 1957; Campbell and Hille, 1976). It may be that lowering external Ca^2+^ could activate more NaP current, contributing to a greater amplification of oscillations. We therefore tested whether changes in Ca^2+^ concentration over the range that modulates oscillation strength would also affect Na^+^ current in fusiform cells. NaP conductance was measured in voltage clamp by gradually ramping membrane potential from -83 to -53 mV over the course of 5 seconds (Fig. 5D; Leao et al. 2012). Voltage control was maintained by limiting Na^+^ current amplitude with a submaximal dose of TTX, by blocking background currents, and by using a narrow range of voltages. In this experiment, we used three Ca^2+^ concentrations, 0.5, 1.7 and 2.5 mM, representing low, moderate and high Ca^2+^, respectively. The amplitude of NaP current at individual membrane voltages was measured and transformed to conductance, which was further plotted as a function of membrane voltage. A Boltzmann function was fitted to the resulting curve in order to calculate the maximum conductance, half-maximal activation voltage (*v50*) and slope factor (see *MATERIALS AND METHODS*). Figure 5E shows normalized conductance curves from a representative neuron to illustrate the difference in *v50*. *V50* displayed a significant positive shift in higher Ca^2+^ concentrations (Fig. 5F, middle; n=9 cells, F(1.7, 13.3)=31.07, p<0.0001, RM one-way ANOVA): *v50* shifted from -61.2 ± 1.0 mV in 0.5 mM to -56.5 ± 1.0 mV in 1.7 mM (p<0.0001) and -56.9 ± 1.3 mV in 2.5 mM (p=0.001). However, *v50* was not measurably different between 1.7 and 2.5 mM conditions (p=0.851). Maximum conductance, as predicted by the Boltzmann fits, was not different between conditions, averaging 3.1 ± 0.7, 3.6 ± 0.8, and 2.7 ± 0.7 nS from low to high concentration (Fig. 5F, left; n=9 cells, F(1.2, 9.9)=1.6, p=0.239, RM one-way ANOVA). Slope was also unchanged between all conditions, averaging 3.0 ± 0.2, 3.2 ± 0.2, and 3.0 ± 0.2 nS/mV in 0.5, 1.7, and 2.5 mM Ca^2+^ (Fig. 5F, right; n=9 cells, F(1.1, 9.0)=0.514, p=0.514, RM one-way ANOVA). To further explore an effect of Ca^2+^ on Na^+^ currents in higher Ca^2+^, in a subset of experiments a wider-range voltage ramp from -83 to -3 mV over the course of 6 seconds was applied to fusiform cells with 1.7 and 2.5 mM Ca^2+^. Consistently, *v50* was not significantly different (1.7 mM Ca^2+^: -56.5 ± 1.4 mV, n=6 cells; 2.5 mM Ca^2+^: -52.8 ± 1.0 mV, n=5 cells; t(9)=2.019, p=0.074, two-tailed t-test). In summary, increasing external Ca^2+^ concentration in the range from 0.5 to 1.7 mM effectively decreased NaP conductance to inhibit oscillations by positively shifting the half-maximal activation voltage of the channels. However, further increases in external Ca^2+^, which further decreased oscillation power, had no measurable effect on the parameters of NaP conductance.

We next investigated what mediates the change in oscillation power between 1.7 and 2.5 mM Ca^2+^. Figure 6A shows the time course of power change from a representative fusiform cell. Similar to what was observed with 0.5 mM Ca^2+^, fusiform cells also responded to 2.5 mM Ca^2+^ in a rapid manner; within three minutes the power dropped below detectable levels, and was restored by re-application of 1.7 mM Ca^2+^. Interestingly, oscillations were preserved when applying 2.5 mM Ca^2+^ along with TEA (Fig. 6A), suggesting that Ca^2+^-activated K^+^ channels might play a role in shunting the voltage change with high external Ca^2+^. In line with this interpretation, input resistance was significantly smaller with 2.5 mM Ca^2+^, but such Ca^2+^ dependence was abolished with 1 mM TEA in the bath (Fig. 6B; average(1.7, 2.5 mM Ca^2+^)= 295.2 ± 42.7, 179.5 ± 26.5 MΩ, n=7 cells, q(6)=6.997, p=0.006; average(1.7+TEA, 2.5 mM Ca^2+^+TEA)=208.1 ± 16.6, 177.1 ± 29.3 MΩ, n=6 cells, q(5)=2.121, p=0.367, Tukey’s post hoc test). Ca^2+^ dependence of oscillation power was also attenuated by TEA or 200 nM apamin to specifically block SK channels. When switching from 1.7 to 2.5 mM Ca^2+^ with TEA or apamin in the bath, oscillations were preserved (Fig. 6C,D). Although we still observed a significant reduction in oscillation power with higher Ca^2+^, the percent reduction is only ∼55% with TEA and ∼56% with apamin, significantly smaller than ∼95% reduction in control (Fig. 6E; control *vs.* TEA, q(16)=12.80, p<0.0001; control *vs.* apamin, q(16)=12.28, p<0.0001, Tukey’s post hoc test). Taken together, elevation of bath Ca^2+^ concentration in the range used in previous brain slice studies activated background SK channel conductance, thereby shunting voltage oscillations.

We also tested for possible effects of Ca^2+^ concentration on HCN conductance (Fig. 7A,B). Ca^2+^ concentration did not affect the maximum HCN tail current (Fig. 7C; average(0.5, 1.7, 2.5 mM Ca^2+^)=245.0 ± 21.5, 254.1 ± 23.8, 265.7 ± 19.7 pA, n=11 cells, F(1.9,19.5)=1.61, p=0.225, RM one-way ANOVA), or the parameters of the Boltzmann equation, including *v50* (Fig. 7C; average(0.5, 1.7, 2.5 mM Ca^2+^)=-92.8 ± 1.4, -92.8 ± 1.2, -92.2 ± 1.4 mV, n=11 cells, F(1.6, 15.9)=0.18, p=0.791, RM one-way ANOVA) and slope (Fig. 7C; average(0.5, 1.7, 2.5 mM Ca^2+^)=-7.6 ± 0.3, -7.5 ± 0.4, -7.5 ± 0.7 pA/mV, n=11 cells, F(1.4, 14.3)=0.03, p=0.929, RM one-way ANOVA). These results indicate that extracellular Ca^2+^ sensitivity of fusiform cell oscillations is attributed to the effect of Ca^2+^ on NaP and SK current, and explains why such slow oscillations in fusiform cells has not been previously reported.

**Figure 7.**
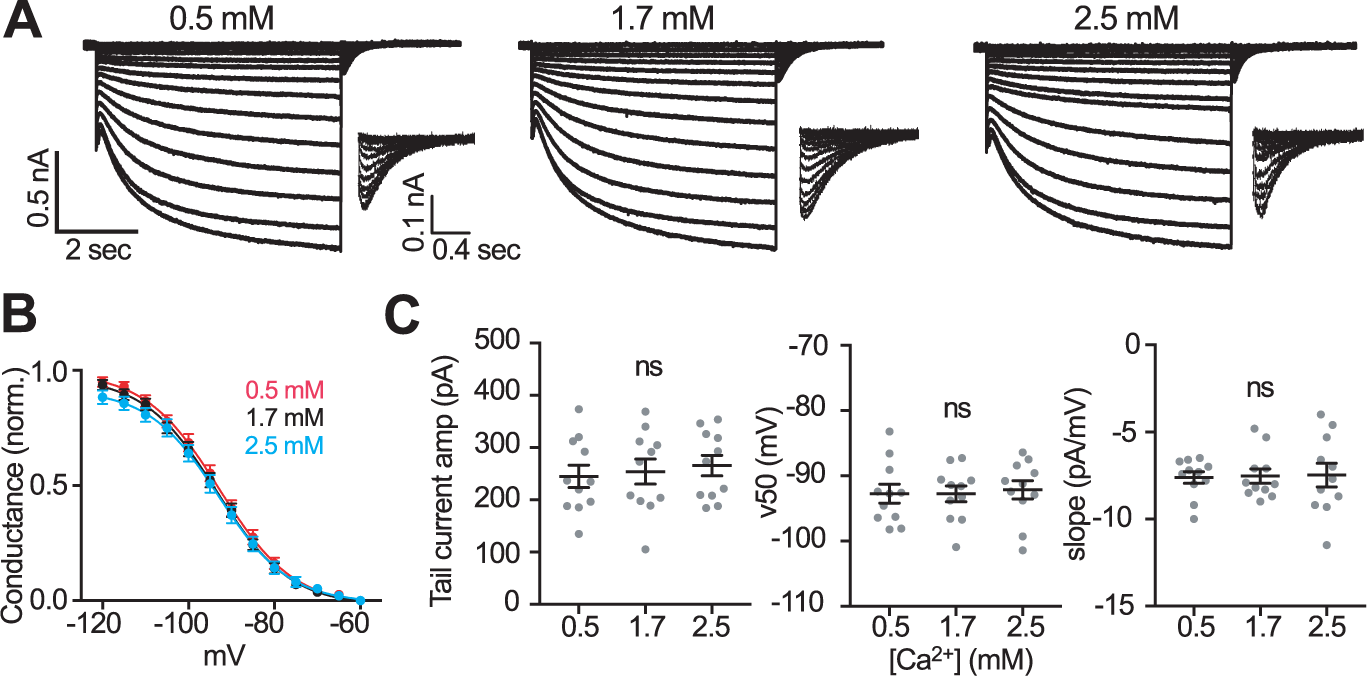
Absence of effect of Ca^2+^ on HCN channel activation. **A**, Example HCN tail currents (magnified in insets) measured in varying concentrations of Ca^2+^: cells were held at -60 mV, injected with current steps from -120 to -60 mV in 5 mV increments for 5 seconds, and stepped back to -60 mV to induce a tail current. **B**, The HCN activation curve was generated by measuring the amplitude of the resulting tail current associated with each voltage step (n=11 cells). Normalized population data are fit with Boltzmann sigmoid equations. **C**, The maximum tail current amplitude, half-activation voltage, and slope did not vary significantly between Ca^2+^ concentrations. Error bar, SEM. Recordings made in the presence of blockers of fast synaptic transmission (5-10 µM NBQX, 10 µM MK-801, 0.5-1 µM strychnine and 5-10 µM SR-95531).

Given the effects of bath Ca^2+^ and of synaptic blockers reported above, we measured the power of oscillations when ongoing synaptic events were present and bath Ca^2+^ held to a physiological level of 1.2 mM. In the absence of synaptic blockers, the power in fusiform cells was significantly larger than that with 1.7 mM Ca^2+^ (Fig. 1F, left; 1.7 Ca^2+^: 0.764 ± 0.122, n=26 cells; 1.2 Ca^2+^: 2.327 ± 0.662 mV^2^/Hz, n=6 cells, U=19.50, p=0.0028, Mann-Whitney U), indicating that oscillations are apparent under physiological conditions.

### Electrical coupling and oscillation synchrony

Fusiform cells are electrically coupled via gap junctions. Although the probability of neighboring fusiform cells being electrically coupled is high (around 70%, based on coupling measured in 20/28 recorded cell pairs), the strength of coupling is low (coupling coefficient ∼0.01; Apostolides & Trussell, 2013). Interestingly, paired recordings from fusiform cells revealed strong oscillation synchrony between neurons despite this apparent weak coupling. Cross-correlograms obtained from cross-correlating the voltage signals of electrically coupled pairs displayed a peak close to zero lag (Fig. 8A). To determine whether the oscillation synchrony was indeed due to electrical coupling, we made dual recordings from Cx36 KO animals in which electrical coupling in DCN is abolished (Apostolides and Trussell, 2013; Yaeger and Trussell, 2016). Interestingly, although fusiform cells from Cx36 KO tissue still displayed oscillations, there was no longer synchrony between closely apposed fusiform cells (Fig. 8B).

**Figure 8.**
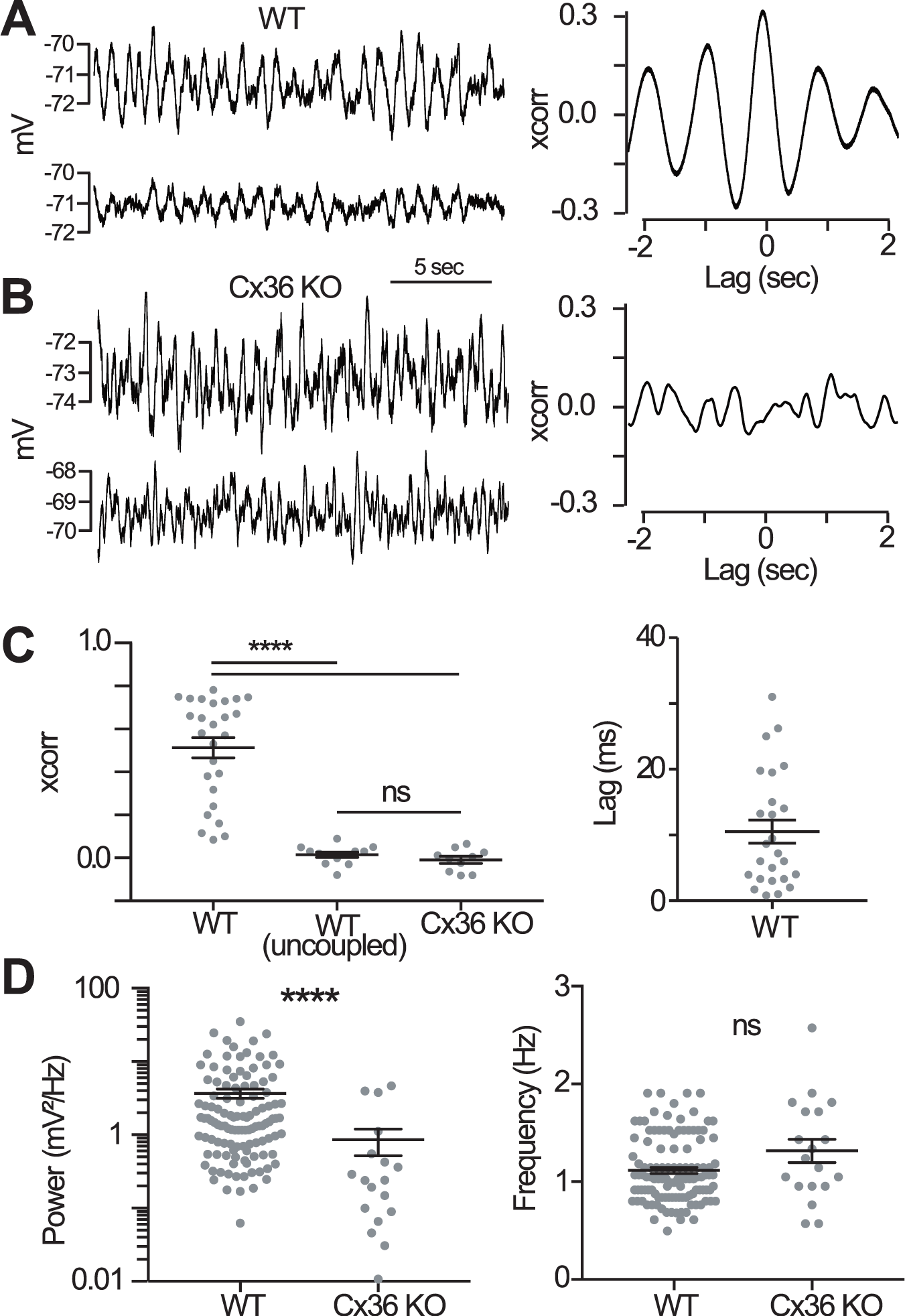
Oscillation synchrony in electrically coupled fusiform cells. **A**, Paired recordings from electrically coupled fusiform cells in WT tissue. The resulting cross-correlogram for this pair, shown to the right, displays a peak around zero lag. **B**, Dual recordings from two fusiform cells in Cx36 KO tissue. The resulting cross-correlogram to the right shows no central peak around zero lag. **C**, *Left*: Electrically coupled cells in WT tissue have significantly larger *xcorr* values (measured at lag zero) than uncoupled pairs in WT and Cx36 KO tissue (F(2,46)=49.6, p<0.0001, one-way ANOVA). *Right*: Lag times measured for the peak in oscillation cross-correlograms for coupled pairs in WT tissue. **D**, Population statistics comparing basic features of oscillations recorded from WT *vs.* Cx36 KO animals. *Left*: Oscillation power was significantly lower in fusiform cells measured from Cx36 KO tissue (U=405, p<0.0001, Mann-Whitney U). *Right*: There was no significant difference in oscillation frequency between WT and KO (U=846, p=0.096, Mann-Whitney U). ****p<0.0001. Error bar, SEM. Recordings made in the presence of blockers of fast synaptic transmission (5-10 µM NBQX, 10 µM MK-801, 0.5-1 µM strychnine and 5-10 µM SR-95531).

Population statistics are displayed in Figure 8C. The average *xcorr* for coupled pairs was 0.5 ± 0.05 (n=26 pairs). This was significantly larger than *xcorr* values for uncoupled pairs in WT and Cx36 KO tissue, both of which were close to zero (Fig. 8C, left; 0.01 ± 0.01 (n=13 pairs) for uncoupled WT and -0.01 ± 0.02 (n=10 pairs) for KO; F(2,46)=49.6, p<0.0001, one-way ANOVA; post hoc WT *vs.* WT uncoupled p<0.0001, WT *vs.* Cx36 KO p<0.0001). Coupled pairs in WT tissue displayed peaks in their cross-correlograms close to zero lag, averaging 10.5 ± 1.8 ms (Fig. 8C, right). These results indicate that fusiform cell oscillations are generated by cell intrinsic conductances, whereas the presence of electrical synapses allows synchrony across the neuronal population (Long et al., 2002).

The oscillations observed in Cx36 KO mice had significantly lower power than oscillations recorded from WT mice (Fig. 8D; averages WT: 3.7 ± 0.5 mV^2^/Hz (n=117 cells), KO: 0.9 ± 0.3 mV^2^/Hz (n=19 cells); U=405, p<0.0001, Mann-Whitney U). The frequency, however, was unchanged (Fig. 8D; averages WT: 1.12 ± 0.03 Hz (n=117 cells), KO: 1.32 ± 0.12 Hz (n=19 cells); U=846, p=0.096, Mann-Whitney U). This result indicates that synchronization rendered by electrical coupling likely acts to boost oscillation power.

### Electrical coupling and spike synchrony

The presence of synchrony in subthreshold voltage oscillations might lead to spike synchrony between fusiform cells (Stefanescu and Shore, 2015, 2017; Wu et al., 2016). To test this possibility, spike times were binned at both 1 and 10 ms and used to generate cross-correlograms of spike times. Fusiform cells were categorized based on their spiking pattern: bursting neurons have >10 spikes per oscillation cycle (Fig. 9A, left), oscillating neurons have <5 spikes per oscillation cycle (Fig. 9B, left), and neurons that do not exhibit oscillatory activity were silent. Electrically coupled neurons in some studies have shown robust spike synchrony, with synchronized spikes occurring within 5 ms of one another (Dugué et al., 2009; Trenholm et al., 2014). By contrast, fusiform pairs showed no evidence of such fine-scale synchrony (Fig. 9A inset), likely due to their relatively weak coupling. However, in pairs where both neurons were exhibiting regular high-frequency spike bursts (Fig. 9A), the resulting cross-correlograms displayed coordinated 1-2 Hz spike oscillations similar to those seen in the voltage cross-correlograms. The average *xcorr* for bursting pairs was 0.22 ± 0.06 (Fig. 9C; n=6 pairs). The oscillations in the cross-correlograms of oscillating pairs with lower frequency spiking (Fig. 9B) were much weaker, resulting in *xcorr* values close to zero (Fig. 9C; oscillating 0.02 ± 0.01, n=6 pairs). The lag time of the peak of these *xcorr* oscillations was similar to lag times measured when correlating oscillating membrane potential, averaging 57.3 ± 12.4 ms (n=11 pairs, compared to 10.5 ± 1.8 ms for voltage oscillation lag). Cross-correlation waveforms for both bursting and oscillating pairs showed an average cycle width of 1012 ± 65 ms (n=11 pairs), matching the subthreshold voltage oscillations of the cells. For each bursting pair, we measured the spike frequency for each cell and averaged them to get the pair spike frequency. We saw a strong positive linear correlation between spike *xcorr* and spike frequency (Fig. 9D; n=12 pairs, R^2^=0.86, p<0.0001, linear regression) (de la Rocha et al., 2007). Increased spike frequency in bursting pairs may therefore increase broad-scale spike synchrony.

**Figure 9.**
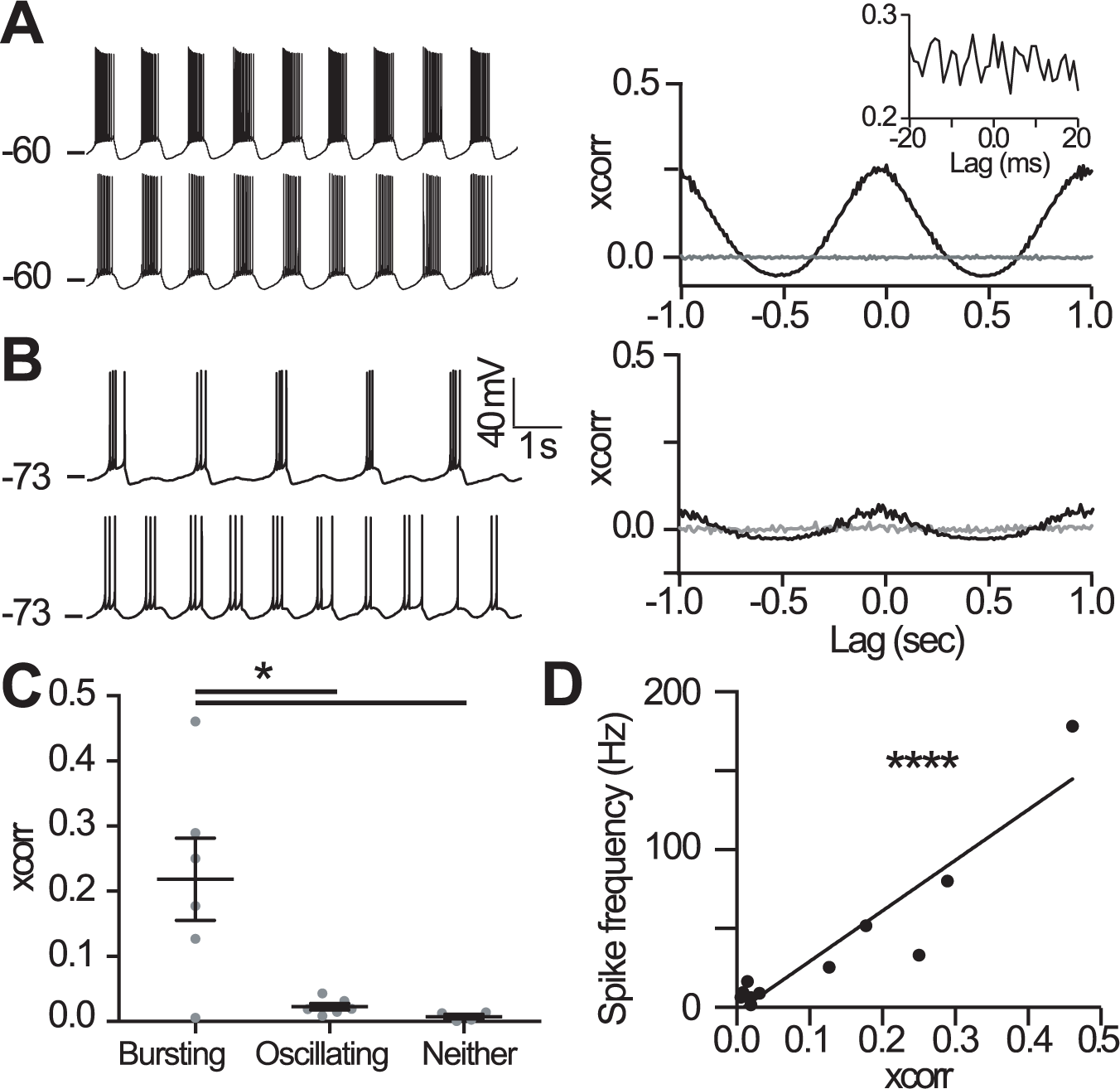
Broad-scale spike synchrony displayed by electrically coupled fusiform cells. **A,** Paired recordings from electrically coupled fusiform cells that are both *bursting* (left) and the resulting spike cross-correlogram for this example pair (right). The inset displays a lack of a peak around zero between ±20 ms: this indicates a lack of fine-scale spike synchrony. Indeed, no pair displayed the fine-scale spike synchrony that is commonly seen in strongly electrically coupled neurons of other brain regions. However, the cross-correlogram on a wider time scale shows a slow oscillation matching the oscillation frequency of the neuron, indicating broad-scale spike synchrony. **B,** Paired recordings from electrically coupled fusiform cells that are both *oscillating* (left) and the resulting spike cross-correlogram for this example pair (right). Pairs either consisted of cells that were bursting (>10 spikes per oscillation cycle), oscillating (<5 spikes per oscillation cycle), or not oscillating. Oscillating pairs showed cross-correlogram oscillations similar to bursting pairs, but much weaker. **C,** *xcorr* measured at zero lag for spike cross-correlograms across the population of fusiform pairs. *xcorr* values measured from bursting pairs were significantly larger than *xcorr* values measured from oscillating and non-oscillating pairs, which both displayed *xcorr* values around 0 (W=8, p=0.01, Kruskal-Wallis). **D,** There was a strong linear correlation between spike *xcorr* and spike frequency in bursting pairs (R^2^=0.86, p<0.0001, linear regression). Spike frequency was averaged between cells to get one value for the pair. *p<0.05, ****p<0.0001. Error bar, SEM. Recordings made in the presence of blockers of fast synaptic transmission (5-10 µM NBQX, 10 µM MK-801, 0.5-1 µM strychnine and 5-10 µM SR-95531).

### Electrical coupling, synchrony, and distance

The distance between somas was approximated for a subset of dual recordings. With the exception of one pair with a distance of 120 μm between cells, all electrically coupled pairs fell within 80 μm of one another (n=9 pairs). There was no significant linear relationship between distance and coupling coefficient (n=20 pairs, R^2^=0.15, p=0.097, linear regression) or oscillation synchrony (n=17 pairs, R^2^=0.19, p=0.119, linear regression). Considering the highly detailed mapping of mouse DCN tonotopy (Muniak et al., 2013) and the relatively conserved region in which paired recordings were made, it is likely that only fusiform cells within the same or similar tonotopic domains were connected.

## DISCUSSION

### Biophysical basis of oscillations

Fusiform cells displayed 1-2 Hz subthreshold voltage oscillations that were synchronized across local clusters of neurons. Given that oscillations were present in synaptic blockers and that the intrinsic resonance frequency was also 1-2 Hz, oscillations and the resonance of fusiform cells are likely determined by the same intrinsic mechanisms (Fransén et al., 2004). Resonance is created by active and passive properties that filter low- and high-frequency voltage fluctuations, respectively, resulting in a peak in the impedance curve (Hutcheon and Yarom, 2000). In fusiform cells, HCN and NaP were required to produce oscillations, consistent with observations made in a variety of other brain regions (Mccormick and Pape, 1990; Fransén et al., 2004; Hu et al., 2009; Boehlen et al., 2013; Matsumoto-Makidono et al., 2016; Stagkourakis et al., 2018). We also observed that electrical connections between fusiform cells boosted oscillation power, indicating that currents generated by NaP and HCN conductances spread through neighboring cells (Apostolides and Trussell, 2014). Modeling studies in other brain regions show that HCN provides a depolarizing drive for the initial rising phase of the oscillation which is further supported or amplified by NaP, whereas continued depolarization deactivates HCN, leading to repolarization (Hutcheon et al., 1996; Dickson et al., 2000; Hutcheon and Yarom, 2000; Fransén et al., 2004). This interplay of voltage-dependent activation and deactivation suggests that mechanisms that control the kinetics of HCN could dictate the basal oscillation frequency in DCN, with NaP regulating its amplitude.

The capacity to generate 1-2 Hz oscillations developed between P9 and P16, and this was paralleled by maturation of other features of fusiform cell firing. Some, such as rheobase and the AP threshold, rate of rise and width, are consistent with a developmental enhancement of Na^+^ current (Benites et al., 2023), thus explaining the lack of oscillations at younger ages. It is notable that hearing in mice begins within this time frame (P12-13), suggesting that sensory activity may be a driving force for electrical maturation of fusiform cells. It will be of interest therefore to examine such electrical development when auditory function is compromised.

Other ion channel blockers had variable effects on oscillations. Although Ca^2+^ channels are required for spontaneous oscillations in inferior olive and lateral olivocochlear neurons (Llinás and Yarom, 1981; Matsumoto-Makidono et al., 2016; Hong et al., 2022), these did not contribute to fusiform cell oscillations. Furthermore, TEA caused an increase in oscillation power, likely by enhancing input resistance. Given that 10 mM TEA blocks low- and high-voltage activated K^+^ channels as well as KCNQ and SK channels (Monaghan et al., 2004; Johnston et al., 2010), these K^+^ channels are unlikely to be involved in generating oscillations, but may instead indirectly regulate oscillation amplitude.

### Effect of external Ca^2+^

External Ca^2+^ had a large effect on oscillation power, likely explaining why previous studies in DCN that used > 2 mM Ca^2+^ had not noted the presence of these rather striking and persistent oscillations (Gardner et al., 2001; Fujino and Oertel, 2003; Manis et al., 2003; Tzounopoulos et al., 2004). Similarly, recent studies (Olsen et al., 2018; Benites et al., 2023) using 2 mM external Ca^2+^ also did not report oscillations. Indeed, lowering external divalent concentrations seems necessary to reveal oscillations in other acute brain slice preparations: Sanchez-Vives & McCormick (2000) observed the emergence of slow oscillations *in vitro* matching those observed *in vivo* when they lowered external divalent concentrations in slice to *in vivo* levels (specifically from 2 to 1 mM Mg^2+^ and from 2 to 1.2 mM Ca^2+^).

The effect of external Ca^2+^ concentration on oscillation power in fusiform cells is two-fold, depending on the range of Ca^2+^ concentration. Between 0.5 mM and 1.7 mM Ca^2+^, oscillation power was affected by a specific action of Ca^2+^ on Na^+^ channels (Campbell and Hille, 1976), because higher external Ca^2+^ induced a significant positive shift in the *v50* for NaP. While an effect of divalents on Na^+^ conductance is generally attributed to charge screening, given that total divalent concentration was held constant, our observations cannot be solely explained by divalent charge screening unless Ca^2+^ is more effective at screening Na^+^ channels than is Mg^2+^ (Horn, 1999). Between 1.7 mM and 2.5 mM, however, Ca^2+^-activated SK channel was a major player that regulates the oscillation strength. Blocking SK channels preserved the oscillations that would otherwise be undectable with 2.5 mM Ca^2+^. Nevertheless, the fact that block of SK channels only preserved ∼50% of oscillation power suggests that other Ca^2+^-sensitive ionic conductances may also modify oscillation strength.

### Synchrony

All electrically-coupled fusiform cell pairs displayed oscillation synchrony. Given the weak coupling strength between fusiform cells, a similar degree of millisecond-precision spike synchrony as seen in some other electrically coupled networks is unlikely (Dugué et al., 2009; Trenholm et al., 2014). Furthermore, fusiform cells are likely coupled in their dendrites, which are far, and thus electrically isolated, from the axon initial segment where spikes are generated (Connors 2017). As such, filter properties of transmission could contribute to the lag in synchrony. By contrast, single spike synchrony between fusiform cells of similar best frequencies is observed *in vivo* (Voigt and Young, 1988; Gochin et al., 1989; Stefanescu and Shore, 2015, 2017). Based on our results, the most likely biological source of such fine-scale spike synchrony *in vivo* is shared synaptic input. Alternatively, given that neuronal pair-wise correlations scale with firing rate even in uncoupled neurons (de la Rocha et al., 2007), the high sound-evoked firing rate of fusiform cells *in vivo* may also promote fine-scale spike synchrony independently of electrical coupling.

### Oscillations in a physiological setting

The DCN processes multisensory inputs and is representative of the class of ‘cerebellum-like’ structures (Oertel and Young, 2004). This kind of structure also includes cerebellar cortex as well as the electrosensory lobe of electric fish. The fact that this unique oscillatory pattern of fusiform cells only emerges after hearing onset suggests that it might play an important role in processing features of auditory signals. Electrical coupling, which boosts oscillation power, appeared in fusiform cells when their somas were within 80 μm of one another. According to frequency maps of mouse DCN, it is likely that only fusiform cells within similar frequency domains are coupled (Muniak et al., 2013). Chemical synapses located close to electrical connections enable synaptic input to spread electrically between cells (Trenholm et al., 2014). NaP and HCN activation by parallel fiber input also reshapes synaptic input, significantly increasing the amplitude and width of the excitatory postsynaptic potential (EPSP), and inducing a damped oscillation following the EPSP (Apostolides and Trussell, 2014). Because gap junctions act as low-pass filters, slower EPSPs are ideal for transmission through gap junctions (Bennett and Zukin, 2004). Furthermore, active dendritic conductances such as NaP amplify synaptically-induced conductances (Schwindt and Crill, 1995). Oscillations would therefore amplify synaptic signals in a synchronized fashion between cells. Thus, electrical coupling may allow for distribution of synaptic input amongst fusiform cells that are within the same frequency domain. Shared variability resulting from synchronized oscillations may work to coordinate the output from a given frequency domain and therefore strengthen the perception of that frequency. Interestingly, similar mechanisms have been proposed in the principal cells of the gymnotiform electrosensory lobe. These neurons also feature slow voltage oscillations that drive spike bursts, as well as amplification of parallel fiber inputs by NaP (Turner et al., 1996; Berman et al., 2001). A proposed function for such oscillations are to provide sensory filters, possibly tuned to accommodate the fish to slow environmental changes or to detect beat frequencies between the fish’s own electrical signals and that of its neighbors. It is possible such oscillatory mechanisms in fusiform cells play a similar role in tuning to environmental or self-generated sensory signals (Singla et al., 2017).

Further study are needed to explore the presence of oscillations in the DCN *in vivo*. Slow oscillations are unlikely to drive spontaneous spike rate as fusiform cells are known to spike at 20-30 Hz *in vivo* (Davis and Young, 2000; Ma and Brenowitz, 2012). Fusiform cells phase-lock to amplitude-modulated tones and have best envelope frequencies, but these modulation frequencies are faster than 1-2 Hz (Kim et al., 1990; Rhode and Greenberg, 1994; Gdowski and Voigt, 1998). The strongest evidence for slow oscillations *in vivo* is associated with spontaneous spike bursts, which in guinea pig occurs at ∼0.2 Hz (Wu et al., 2016). HCN/NaP oscillations could underlie these burst rates, particularly if more intact electrical coupling *in vivo* imposes a slower network frequency (Stagkourakis et al., 2018). Alternatively, if the conductance state of fusiform cells is higher *in vivo* than in brain slices (Fernandez et al., 2018), slow oscillations in fusiform cells may effectively be compartmentalized to dendrites and thus exert their function locally. Indeed, the distal apical dendrites of fusiform cells are likely where electrical coupling occurs, given that this is the site where they are also coupled with molecular-layer interneurons (Apostolides and Trussell, 2013).

## Acknowledgements

This work was supported by National Institutes of Health Grants DC004450 to L.O.T. and DC015187 to L.A.M. We thank Dr. Gideon Rothschild and members of the Trussell laboratory for helpful discussions and Sean Elkins, Jennifer Goldsmith, Ruby Larisch, and Michael Bateschell for assistance with mouse colony management. Current address for LAM is Masonic Institute for the Developing Brain, Institute of Child Development, College of Education and Human Development, University of Minnesota, Minneapolis, MN 55414. Current address for PFA is Kresge Hearing Research Institute & Department of Otolaryngology, University of Michigan, Ann Arbor, MI 48109.

---

**Extended Data - Table 2-1.**
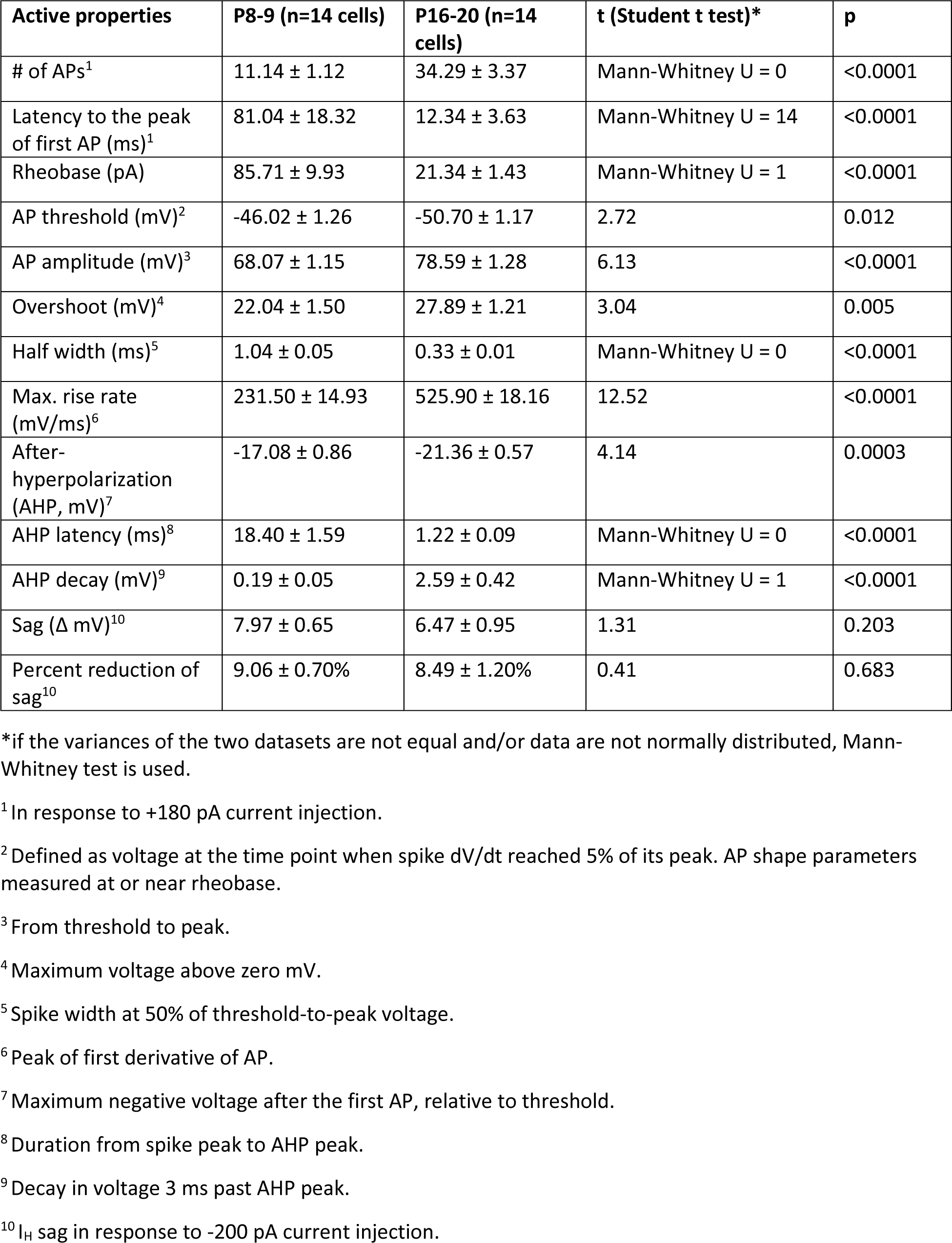
Development of active properties in fusiform cells (mean ± SEM)

| Active properties | P8-9 (n=14 cells) | P16-20 (n=14 cells) | t (Student t test)* | p |
| --- | --- | --- | --- | --- |
| # of APs<sup>1</sup> | 11.14 ± 1.12 | 34.29 ± 3.37 | Mann-Whitney U = 0 | <0.0001 |
| Latency to the peak of first AP (ms)<sup>1</sup> | 81.04 ± 18.32 | 12.34 ± 3.63 | Mann-Whitney U = 14 | <0.0001 |
| Rheobase (pA) | 85.71 ± 9.93 | 21.34 ± 1.43 | Mann-Whitney U = 1 | <0.0001 |
| AP threshold (mV)<sup>2</sup> | -46.02 ± 1.26 | -50.70 ± 1.17 | 2.72 | 0.012 |
| AP amplitude (mV)<sup>3</sup> | 68.07 ± 1.15 | 78.59 ± 1.28 | 6.13 | <0.0001 |
| Overshoot (mV)<sup>4</sup> | 22.04 ± 1.50 | 27.89 ± 1.21 | 3.04 | 0.005 |
| Half width (ms)<sup>5</sup> | 1.04 ± 0.05 | 0.33 ± 0.01 | Mann-Whitney U = 0 | <0.0001 |
| Max. rise rate (mV/ms)<sup>6</sup> | 231.50 ± 14.93 | 525.90 ± 18.16 | 12.52 | <0.0001 |
| After-hyperpolarization (AHP, mV)<sup>7</sup> | -17.08 ± 0.86 | -21.36 ± 0.57 | 4.14 | 0.0003 |
| AHP latency (ms)<sup>8</sup> | 18.40 ± 1.59 | 1.22 ± 0.09 | Mann-Whitney U = 0 | <0.0001 |
| AHP decay (mV)<sup>9</sup> | 0.19 ± 0.05 | 2.59 ± 0.42 | Mann-Whitney U = 1 | <0.0001 |
| Sag ($\Delta$ mV)<sup>10</sup> | 7.97 ± 0.65 | 6.47 ± 0.95 | 1.31 | 0.203 |
| Percent reduction of sag<sup>10</sup> | 9.06 ± 0.70% | 8.49 ± 1.20% | 0.41 | 0.683 |

\*if the variances of the two datasets are not equal and/or data are not normally distributed, Mann-Whitney test is used.
<sup>1</sup> In response to +180 pA current injection.
<sup>2</sup> Defined as voltage at the time point when spike dV/dt reached 5% of its peak. AP shape parameters measured at or near rheobase.
<sup>3</sup> From threshold to peak.
<sup>4</sup> Maximum voltage above zero mV.
<sup>5</sup> Spike width at 50% of threshold-to-peak voltage.
<sup>6</sup> Peak of first derivative of AP.
<sup>7</sup> Maximum negative voltage after the first AP, relative to threshold.
<sup>8</sup> Duration from spike peak to AHP peak.
<sup>9</sup> Decay in voltage 3 ms past AHP peak.
<sup>10</sup> $I_H$ sag in response to -200 pA current injection.

